# Reverse Transcriptase Real-Time PCR Assay for the Rapid Enumeration of Live *Candida auris* from the Healthcare Environment

**DOI:** 10.1101/2021.03.18.436063

**Authors:** Bryanna Lexus Freitas, Lynn Leach, Vishnu Chaturvedi, Sudha Chaturvedi

**Author notes:** Corresponding Author: Sudha Chaturvedi. Bryanna Freitas and Lynn Leach contributed equally to the study. Bryanna Freitas was assigned first author as she devoted fulltime laboratory effort to this project and performed key experiments while learning many techniques. Lynn Leach assembled key reagents/samples for PCR, trained Bryanna Freitas on PCR assays, and performed few crucial experiments independently and in conjunction with Bryanna Freitas.

## Abstract

Ongoing healthcare-associated outbreaks of multidrug-resistant yeast *Candida auris* have prompted development of several rapid DNA-based molecular diagnostic tests. These tests do not distinguish between live and dead *C. auris* cells, limiting their use for environmental surveillance and containment efforts. We addressed this critical gap by developing a reverse transcriptase (RT) real-time PCR assay for rapid detection of live *C. auris* in healthcare environments. The assay targets the internal transcribed spacer 2 (ITS2) cDNA of *C. auris*, amplified by obtaining pure RNA followed by reverse transcription and real-time PCR assay. The assay was highly sensitive, with the detection limit of ten colony-forming units (CFU) per RT real-time PCR reaction. In *C. auris* viability studies, ITS2 cDNA was detectable in heat- and ethanol-killed *C. auris* cells, but not bleach-killed cells. Validation studies showed no cycle threshold (Ct) values were obtained from samples on a sponge matrix spiked with either dead *C. auris* (10^5^/ml) or other *Candida* species (10^5^/ml), while most environmental samples spiked with 10^2^ to 10^5^ CFU of live *C. auris* yielded positive Ct values. Finally, 33 environmental samples positive for *C. auris* DNA but negative by culture were all negative by RT real-time PCR assay, confirming concordance between culture and the assay. The RT real-time PCR assay appears highly reproducible, robust, and specific for detecting live *C. auris* from environmental samples, and could be an invaluable tool in surveillance efforts to control the spread of live *C. auris* in healthcare environments.

## INTRODUCTION

*Candida auris* is an emerging pathogenic yeast of significant clinical concern because of its frequent intrinsic resistance to fluconazole and other antifungal drugs, and the high mortality associated with systemic infections (1-3). *C. auris* infections emerged in the United States in 2016, and New York state faced an unprecedented outbreak with 50-60% mortality in patients with invasive infection (2, 4, 5). *C. auris* South Asia Clade I was linked to the New York outbreak (5), which is particularly concerning as some strains of South Asia Clade I are now resistant to all three antibiotic classes used to treat fungal infection (6, 7). *Candida auris* has frequently been recovered from environmental surfaces in rooms of colonized or infected patients (4), and a report of an outbreak of *C. auris* in an intensive care unit linked transmission to shared temperature probes (3). The ability of *C. auris* strains to survive for prolonged periods on dry and moist surfaces is documented (8). Recent studies showed heavy colonization of skin and mucous membranes of hospitalized patients as well as various healthcare objects by *C. auris*, suggesting that both patients and contaminated surfaces might be a source of *C. auris* transmission in the healthcare setting (5, 9). In 2019, the Centers for Disease Control and Prevention (CDC) added *C. auris* to its list of ‘urgent threats’ to public health apart from four drug-resistant bacterial pathogens (https://www.cdc.gov/drugresistance/pdf/threats-report/candida-auris-508.pdf). Given the importance of *C. auris* as a healthcare-associated pathogen, it is concerning that the environmental reservoirs for this pathogen are currently not known. *C. auris* exhibits relative resistance to widely used quaternary ammonium disinfectants (10) and thus far, only bleach products have been shown to be effective in eradicating *C. auris* from the environment (11).

Rapid testing methods are needed for the detection and prevention of this deadly fungal pathogen. In the last several years, extensive work was carried out to identify *C. auris* from patient samples. These included utilization of selective medium for the recovery of *C. auris* in culture (12), as well as optimization of biochemical (https://www.cdc.gov/fungal/candida-auris/pdf/Testing-algorithm_by-Method_508.pdf), and protein-based databases (http://www.cidrap.umn.edu/news-perspective/2018/04/fda-approves-rapid-diagnostic-test-candida-auris & https://www.rapidmicrobiology.com/news/new-fda-clearance-for-vitek-ms-expanded-id-for-challenging-pathogens) for *C. auris* identification. Though these methods are excellent, they take considerable time (a few days to weeks) to confirm identification of *C. auris* (5). To combat these issues, we and others have developed molecular-based assays using various real-time PCR chemistries (13, 14, 15, 16). All molecular diagnostics developed thus far for *C. auris* have relied on DNA, which does not differentiate live from dead *C. auris* cells. This drawback has hampered environmental surveillance and containment efforts as the culturing of surveillance samples followed by identification can take anywhere from 4-14 days.

The detection of RNA by reverse transcription (RT) PCR has shown promise for the detection of viable microorganisms (17-19). Therefore, we used primers/probe previously designed from internal transcribed spacer 2 (*ITS2*) (13) to obtain its reverse transcription (ITS2 cDNA) followed by real-time PCR. The suitability of ITS2 cDNA as the indicator of *C. auris* viability was validated on environmental surveillance samples. The excellent concordance between live *C. auris* in culture and positive Ct values through RT real-time PCR was observed. This new RT real-time PCR assay could be an invaluable tool in surveillance efforts to control the spread of live *C. auris* in the healthcare environment.

## MATERIAL AND METHODS

### Yeast Strains and Media

*C. auris* isolate M5658, a major genotype belonging to the South Asia Clade I and involved in the current outbreak in New York, was used for all standardization experiments of the reverse transcriptase real-time PCR assay. Additionally, *C. auris* strains from East Asia Clade II, South Africa Clade III, South America Clade IV and isolates of *C. albicans, C. glabrata. C. parapsilosis, C. tropicalis, C. haemulonii*, and *C. duobushhaemulonii* were used for assay specificity (Supplementary Table 1). Sabouraud dextrose agar (SDA) with or without chloramphenicol (25 mg/liter) was used for the culturing of all *Candida* spp.

### RNA extraction

*C. auris* cells, grown on SDA for 48-72 h, were scrapped with a sterile inoculation loop, and cell suspension was prepared in phosphate buffered saline (PBS) containing 0.1% bovine serum albumin (BSA). Cells were washed twice with PBS-BSA, resuspended in same buffer, and OD_530_ = 0.1 was determined. This measurement resulted in 10^6^ CFU/ml on SDA plate (Anower et al., personal communication). Serial 10-fold dilutions of *C. auris* cell suspension prepared in PBS-BSA were centrifuged at 12,500 RPM for 1 min and RNA from pellet was extracted using MasterPure Yeast RNA Purification kit (Epicenter®). In brief, each pellet was suspended in 300 μl of extraction reagent containing 1 μl of 50 μg/μl Proteinase K and mixed with a vortex shaker (1000 rpm/min). The mixture was incubated at 70 oC for 15 min in the water bath with intermittent mixing every 5 min. Following incubation, samples were placed in ice for 5 min, and then 175 μl of MPC Protein Precipitation Reagent was added to 300 μl of lysed samples. The mixture was vortexed briefly for 10 sec and then centrifuged at 12,500 RPM for 10 min at 4oC. Supernatant was transferred to a clean tube and the pellet was discarded. The RNA in the supernatant was precipitated by adding 500 μl of isopropanol, inverting the tube 30-40 times and centrifuging at 12,500 RPM at 4oC. Isopropanol was removed without dislodging the pellet. The pellet was washed twice with 70% ethanol and RNA was resuspended in 40 μl of Tris EDTA (TE) buffer. Any DNA contamination in the extracted RNA was removed by treating each sample with 5 μl of TURBO DNAase buffer and 1 μl of TURBO DNase enzyme Thermo Fisher Scientific). Samples were mixed gently by flicking the tube followed by incubation at 37oC for 30 min in the water bath. Following incubation, 5 μl of DNase Inactivation Reagent was added, and samples were incubated at room temperature for 5 min with intermittent flicking. The samples were centrifuged at 12,500 RPM for 5 min at room temperature, then liquid was transferred to a new tube and the pellet was discarded. All purified RNA was stored at −20°C until further testing. RNA from other *Candida* species was also extracted using the same procedure.

RNA was also extracted from environmental samples (sponge) collected from various health care facilities. Each sample was placed in a Whirl-Pak Homogenizer Blender Filter bag containing 45•ml of PBS containing 0.02% Tween 80 (PBS-T80). The bags were gently mixed in the Stomacher 400 Circulator (Laboratory Supply Network, Inc., Atkinson, NH, USA) at 260•rpm for 1•min to remove the content of the sponge in the liquid, The liquid was transferred into a 50-ml conical tube and centrifuged at 4,000•rpm for 5•min, and the supernatant was decanted, leaving about 3•ml of liquid at the bottom of the tube. The 3•ml liquid was vortexed briefly, and from this, 1•ml aliquots of surveillance samples were drawn and spiked with 10-fold dilution series of *C. auris*, then RNA was extracted as described above. RNA was also extracted from un-spiked environmental samples.

### Quantitative Reverse Transcriptase Real-Time PCR Assay

The Ultra Plex 1-Step Tough Mix (Quanta Bio), which is a ready-to-use 4X concentrated master mix for reverse transcription (RT) real-time PCR of RNA templates using hybridization probe detection chemistries (e.g., TaqMan® 5’-hydrolysis probes) on ABI7500 (Applied Biosystems) was used. In this system, the first-strand cDNA synthesis and PCR amplification were carried out in the same tube without opening between the procedures. The PCR amplification was carried out in 20 μl final volume containing 5 μl of RNA template, 1 μl (500 nM) each of ITS2 forward (V2424F (CAURF), 5•-CAGACGTGAATCATCGAATCT-3), and reverse V2426 (CAURR), 5•-TTTCGTGCAAGCTGTAATTT-3) primers, 0.8 μl (100 nM) of ITS2 (V2425P (CAURP), 5•-/56-carboxyfluorescein (FAM)/AATCTTCGC/ZEN/GGTGGCGTTGCATTCA/3IABkFQ/-3•) probe, 5 μl of Ultra Plex 1-Step Tough Mix, and 7.2 μl of nucleic acid free water. All PCR run also included 5 μl of positive extraction (*C. auris* M5658; 10^3^ CFU/50 •l) and positive amplification (*C. auris* M5658, 0.02 pg/μl), 5 μl of negative extraction (reagents only) and negative amplification (nuclease free water) control. All experiments with internal positive control (VetMAX™ XENO™) were prepared by adding 1.2 μl of XENO control to the master mix and the volume of water in the master mix was reduced to 6.0 μl. The probe in the internal control was tagged with VIC dye.

Reverse transcription and real-time PCR reactions were carried out in the ABI 7500 FAST (Applied Biosystems) instrument starting with cDNA synthesis at 50°C for 10 min, followed by initial denaturation at 95°C for 3 min and real-time PCR cycling (45 cycles) conditions of denaturation at 95°C for 3 sec and annealing at 60°C for 30 sec. The cycle threshold (Ct) was determined by the instrument. The assay sensitivity and specificity were analyzed in triplicate, while assay reproducibility and retrospective studies were done in duplicate. The coefficient of efficiency (*E*) was calculated using the formula E = (10-^1/slope^)-1. Ct values of ≤40 were considered positive and Ct values of >40 were considered negative. All RNA extracted samples were also assessed for DNA contamination by real-time PCR assay as described previously (13).

### Reverse Transcriptase Real-Time PCR Assay Sensitivity, Reproducibility, and Specificity

*C. auris* isolate M5658, a major genotype belonging to the South Asia clade I and involved in the current outbreak in New York, was used to assess the analytical sensitivity of the RT real-time PCR assay. The analytical sensitivity was determined in 10-fold serial dilutions of *C. auris* cells in PBS-BSA and in environmental sponge matrix. In brief, environmental sponge samples negative for *C. auris* DNA by earlier established real-time PCR assay (13) and also negative for *C. auris* by culture were pooled and spiked with various concentration of *C. auris* cells. All spiked samples were processed for RNA extraction and five microliters of the extracted RNA were used in the RT real-time PCR assay in triplicate.

Since primers and probe were assessed extensively for assay specificity in our previous investigation (13), only a small panel of *Candida* spp. were tested in the present investigation. In brief, 5 μl of extracted RNA from each of the *C. auris* clades I to IV and other *Candida* spp., including *C. haemulonii, C. duobushhaemulonii, C. albicans, C. glabrata, C. parapsilosis* and *C. tropicalis*, was run in RT real-time PCR assay in triplicate. The assay reproducibility was assessed by spiking pooled negative surveillance samples with various concentrations of live *C. auris*, bleach-killed *C. auris* and other *Candida* spp., followed by RNA extraction and RT real-time PCR assay.

### Retrospective Analysis of Environmental Surveillance Samples

A total of 33 surveillance samples collected in the year 2020, stored at 4°C, and with 24 of the 33 positive by earlier real-time PCR assay (13), were part of this investigation. A one-ml aliquot of each sample was processed for RNA extraction as described above, followed by RT real-time PCR assay.

### Heat, Bleach, and ethanol Inactivation of *C. auris*

*C. auris* (M5658) grown on SDA plate for 72 h were scrapped and suspended in PBS-BSA and OD_530_ = 0.1 (equivalent to 10^6^ CFU/ml) was determined. Several 1 ml aliquots were incubated at 90 °C for 1 h with shaking at 400 RPM in the Eppendorf ThermoMixer® C. RNA was extracted from the sample just after incubation, and then 30 min, 1 h, 2 h, 4 h, 24 h, and 48 h post-incubation at room temperature followed by RNA extraction and RT real-time PCR assay. For bleach treatment, each 1 ml aliquot of *C. auris* cells was washed with PBS-BSA, resuspended in 900 μl of PBS-BSA, and 100 μl of bleach from a stock bottle was added to get a final concentration of 10% bleach per tube. For ethanol treatment, each *C. auris* cell suspension aliquot was washed with PBS-BSA and resuspended in 300 μl of PBS-BSA. To get a final concentration of 70% ethanol, 700 μl of 100% ethanol was added to each tube. Bleach addition and ethanol treatment were carried out at room temperature for 1, 3, 5, 10, and 30 min followed by washing with PBS-BSA, RNA extraction and RT real-time PCR assay. Another set of *C. auris* cells were treated with 70% ethanol for 10 min followed by washing cells with PBS-BSA and incubation at room temperature for 30 min to 48 h. Cell viability of each treatment was assessed by plating *C. auris* cell suspension (10^6^ CFU/ml) on SDA plate and incubating at 30 °C for 96 h.

### Statistical analysis

The results were statistically analyzed using GraphPad Prism 8.0 Software for macOS. The statistical degree of significance was set at a P value of < 0.05. All the Ct values were averages of at least three repetitions for sensitivity, specificity and cellular viability experiments.

## RESULTS

### Reverse Transcriptase real-time PCR (RT-rt-PCR) Assay Sensitivity, Specificity, and Reproducibility

The RT real-time PCR assay was linear over 5 orders of magnitude, and the limit of detection of the assay was 10 CFU/PCR reaction in PBS-BSA and sponge matrix, respectively (Fig. 1). The assay was highly specific, as no PCR amplification was detected for *Candida* species other than *C. auris* (Supplementary Table 1). The assay was highly reproducible when blinded surveillance samples spiked with either live or bleach-killed *C. auris* or live *Candida* spp. other than *C. auris* were analyzed. Of 17 environmental samples spiked with low to high numbers of live *C. auris* cells (10^2^ to 10^5^), 15 yielded positive results for *C. auris* ITS2 cDNA by RT real-time PCR, while all un-spiked as well as spiked-environmental samples with bleach-killed *C. auris* and other live *Candida* spp. were negative, confirming assay reproducibility and specificity (Table 1A). Using the culture method as the gold standard, the accuracy of the RT real-time PCR assay for the detection of live *C. auris* from surveillance sample was 98%, with positive predictive value and negative predictive value of 100% and 87%, respectively (Table 1B).

**Table 1A.** *C. auris* RT real-time PCR assay reproducibility using blinded panels.

| Samples<br>No. | Yeast Spp.<br>Spiked in<br>Surveillance<br>Sample | No.<br>Yeast<br>Cells<br>Spiked<br>per<br>Sample | No. Yeast<br>cells/PCR<br>Rxn | Live<br>or<br>Dead<br>Yeast | RT real-time PCR |  |  | Internal<br>Control<br>Mean<br>Ct | Final<br>Interpretation |
| --- | --- | --- | --- | --- | --- | --- | --- | --- | --- |
|  |  |  |  |  | Ct 1 | Ct 2 | Mean<br>Ct |  |  |
| 1 | *C. albicans* | 10<sup>5</sup> | 10<sup>4</sup> | Live | 0.0 | 0.0 | 0.0 | 32.26 | NEGATIVE |
| 2 | *C. auris* | 10<sup>5</sup> | 10<sup>4</sup> | Live | 23.26 | 23.89 | 23.62 | 32.42 | POSITIVE |
| 3 | *C. auris* | 10<sup>5</sup> | 10<sup>4</sup> | Live | 23.31 | 23.99 | 23.65 | 32.81 | POSITIVE |
| 4 | *C. auris* | 10<sup>4</sup> | 10<sup>3</sup> | Live | 28.32 | 28.11 | 28.21 | 32.36 | POSITIVE |
| 5 | *C. auris* | 10<sup>3</sup> | 10<sup>2</sup> | Live | 33.17 | 32.96 | 33.06 | 32.42 | POSITIVE |
| 6 | *C. albicans* | 10<sup>5</sup> | 10<sup>4</sup> | Live | 0.0 | 0.0 | 0.0 | 32.52 | NEGATIVE |
| 7 | *C. auris* | 10<sup>3</sup> | 10<sup>2</sup> | Live | 32.77 | 28.9 | 30.83 | 33.41 | POSITIVE |
| 8 | None | - |  | - | 0.0 | 0.0 | 0.0 | 32.58 | NEGATIVE |
| 9 | *C. auris* | 10<sup>4</sup> | 10<sup>3</sup> | Live | 27.08 | 28.9 | 28.0 | 32.45 | POSITIVE |
| 10 | *C. auris* | 10<sup>5</sup> | 10<sup>4</sup> | Dead | 0.0 | 0.0 | 0.0 | 32.44 | NEGATIVE |
| 11 | *C. auris* | 10<sup>5</sup> | 10<sup>4</sup> | Dead | 0.0 | 0.0 | 0.0 | 32.93 | NEGATIVE |
| 12 | *C. auris* | 10<sup>4</sup> | 10<sup>4</sup> | Dead | 0.0 | 0.0 | 0.0 | 32.02 | NEGATIVE |
| 13 | None | - | - | - | 0.0 | 0.0 | 0.0 | 32.33 | NEGATIVE |
| 14 | *C. glabrata* | 10<sup>5</sup> | 10<sup>4</sup> | Live | 0.0 | 0.0 | 0.0 | 32.55 | NEGATIVE |
| 15 | *C. auris* | 10<sup>4</sup> | 10<sup>4</sup> | Dead | 0.0 | 0.0 | 0.0 | 33.33 | NEGATIVE |
| 16 | *C. auris* | 10<sup>2</sup> | 10<sup>1</sup> | Live | 35.74 | 37.62 | 36.68 | 33.98 | POSITIVE |
| 17 | *C. auris* | 10<sup>5</sup> | 10<sup>4</sup> | Live | 24.64 | 24.93 | 24.75 | 32.40 | POSITIVE |
| 18 | *C. auris* | 10<sup>4</sup> | 10<sup>3</sup> | Live | 28.63 | 24.94 | 26.78 | 32.12 | POSITIVE |
| 19 | *C. auris* | 10<sup>2</sup> | 10<sup>1</sup> | Live | 37.98 | 37.00 | 37.48 | 33.20 | POSITIVE |
| 20 | *C. auris* | 10<sup>2</sup> | 10<sup>1</sup> | Live | 0.0 | 0.0 | 0.0 | 32.76 | NEGATIVE<sup>s</sup> |
| 21 | *C. auris* | 10<sup>2</sup> | 10<sup>1</sup> | Live | 38.0 | 40.72 | 39.32 | 32.76 | POSITIVE |
| 22 | *C. auris* | 10<sup>2</sup> | 10<sup>1</sup> | Live | 0.0 | 0.0 | 0.0 | 33.15 | NEGATIVE<sup>s</sup> |
| 23 | None | - | - | - | 0.0 | 0.0 | 0.0 | 33.87 | NEGATIVE |
| 24 | *C. auris* | 10<sup>5</sup> | 10<sup>4</sup> | Live | 24.71 | 25.5 | 25.10 | 32.42 | POSITIVE |
| 25 | *C. auris* | 10<sup>4</sup> | 10<sup>3</sup> | Live | 26.99 | 25.59 | 26.29 | 32.98 | POSITIVE |
| 26 | *C. haemulonii* | 10<sup>5</sup> | 10<sup>4</sup> | Live | 0.0 | 0.0 | 0.0 | 32.76 | NEGATIVE |
| 27 | *C. duobushaemolunii* | 10<sup>5</sup> | 10<sup>4</sup> | Live | 0.0 | 0.0 | 0.0 | 32.75 | NEGATIVE |
| 28 | *C. haemulonii* | 10<sup>5</sup> | 10<sup>4</sup> | Live | 0.0 | 0.0 | 0.0 | 34.40 | NEGATIVE |
| 29 | *C. auris* | 10<sup>5</sup> | 10<sup>4</sup> | Live | 22.5 | 22.3 | 22.4 | 32.62 | POSITIVE |
| 30 | *C. auris* | 10<sup>5</sup> | 10<sup>4</sup> | Live | 22.94 | 22.76 | 22.85 | 33.24 | POSITIVE |
| NEC | - | - | - | - | 0.0 | 0.0 | 0.0 | - | NEGATIVE |
| XENO | - | - | - | - | - | - | - | 31.57 | POSITIVE |
| NTC | - | - |  | - | 0.0 | 0.0 | 0.0 | - | NEGATIVE |
| PAC | *C. auris* | 10<sup>4</sup> | 10<sup>3</sup> | Live | 28.01 | 28.16 | 28.08 | - | POSITIVE |

\$ Samples showing discrepant results

**Table 1B.**
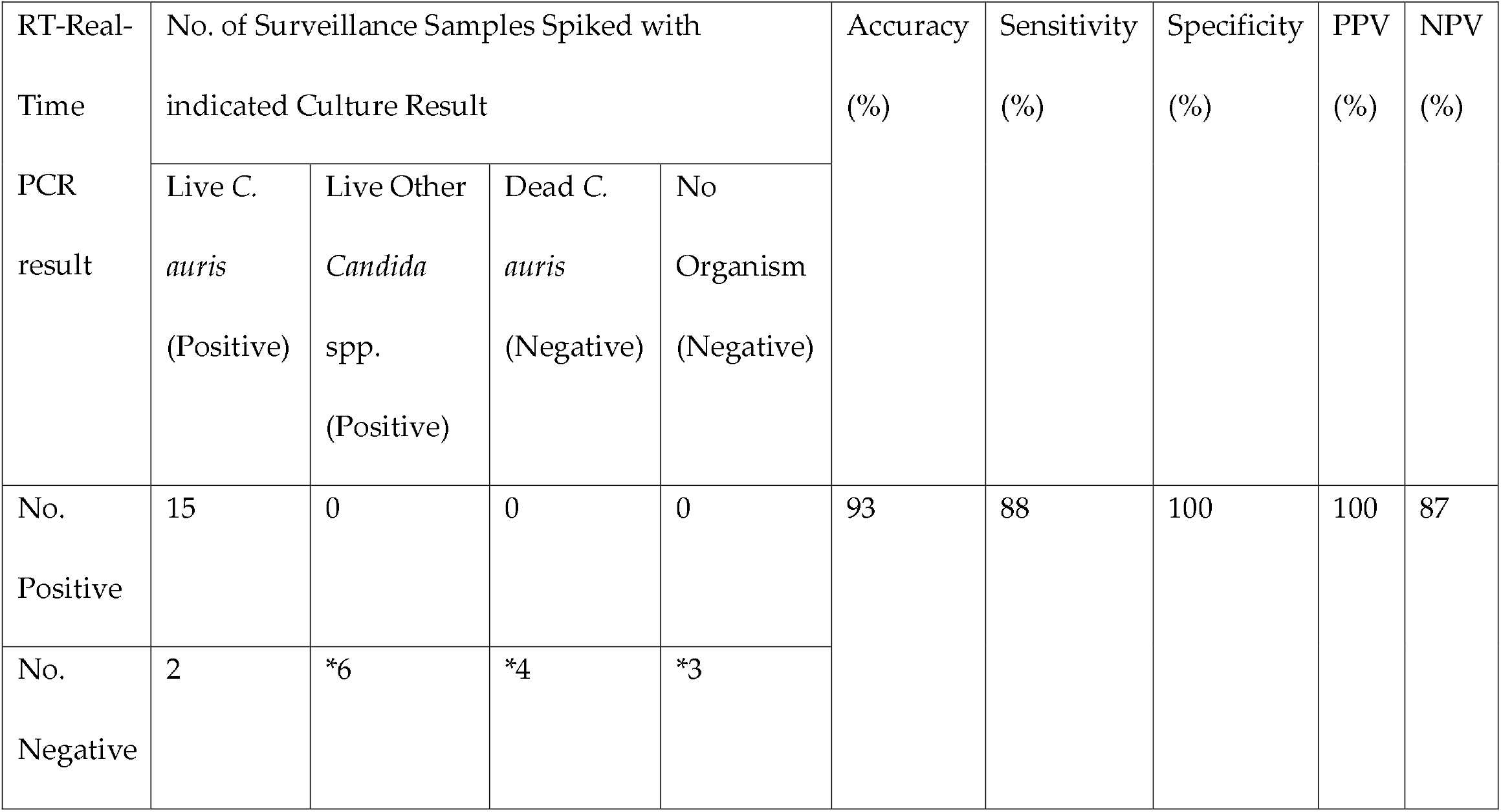
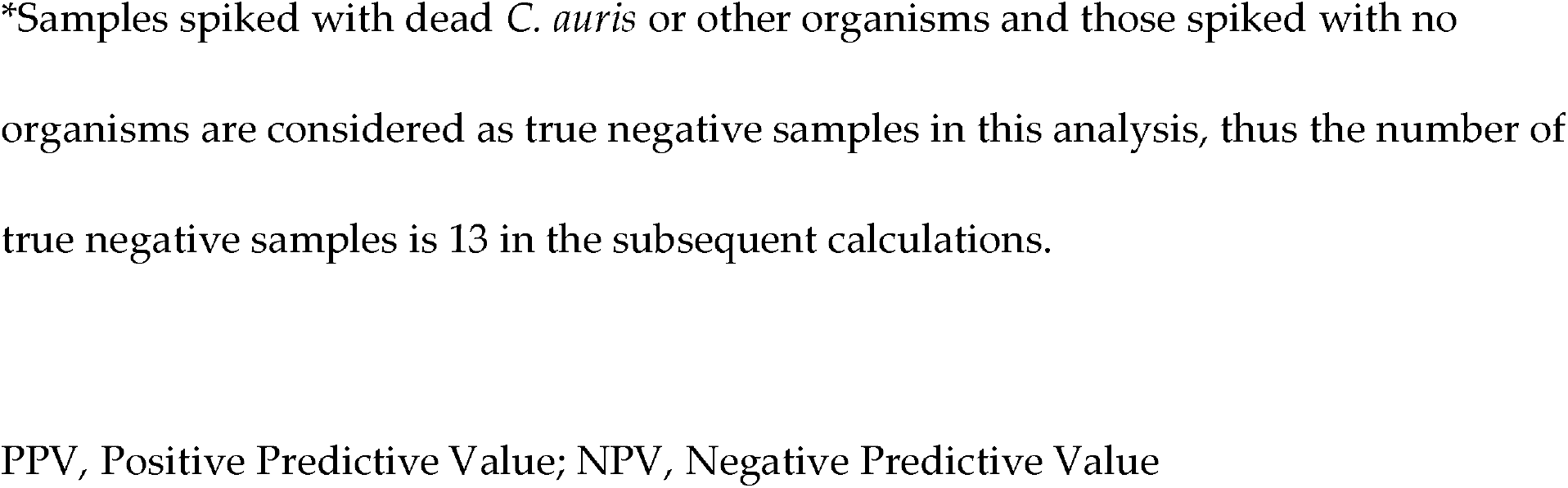
Comparison of culture and RT real-time PCR assay of spiked blinded surveillance samples.

| RT-Real-Time PCR result | No. of Surveillance Samples Spiked with indicated Culture Result |  |  |  | Accuracy (%) | Sensitivity (%) | Specificity (%) | PPV (%) | NPV (%) |
| --- | --- | --- | --- | --- | --- | --- | --- | --- | --- |
|  | Live *C. auris* (Positive) | Live Other *Candida* spp. (Positive) | Dead *C. auris* (Negative) | No Organism (Negative) |  |  |  |  |  |
| No. Positive | 15 | 0 | 0 | 0 | 93 | 88 | 100 | 100 | 87 |
| No. Negative | 2 | *6 | *4 | *3 |  |  |  |  |  |

\*Samples spiked with dead *C. auris* or other organisms and those spiked with no organisms are considered as true negative samples in this analysis, thus the number of true negative samples is 13 in the subsequent calculations.
PPV, Positive Predictive Value; NPV, Negative Predictive Value

**Fig. 1.**
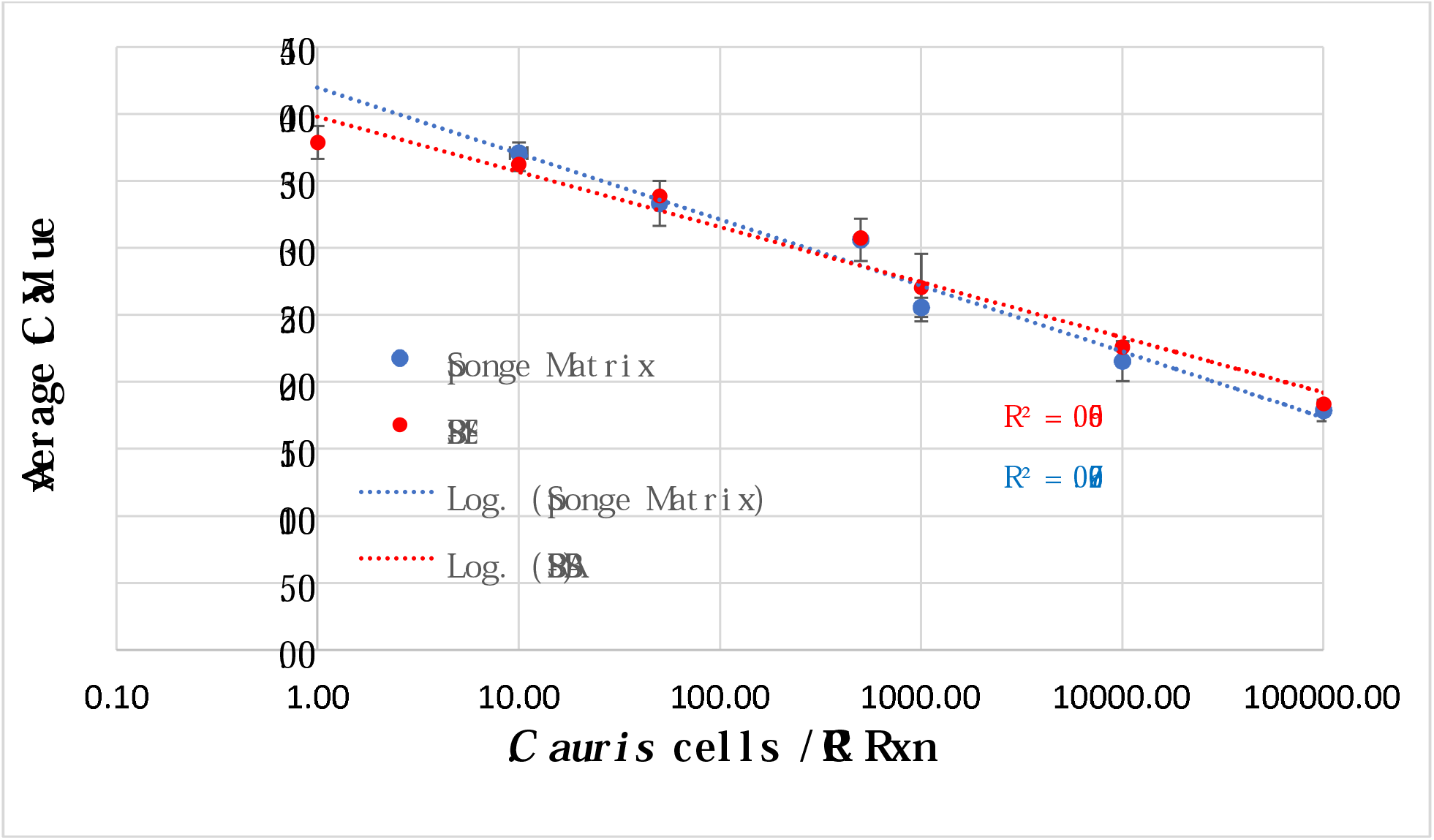
*C. auris* RT real-time PCR assay sensitivity.

### Stability of ITS2 RNA in Heat, 70% ethanol, and Bleach-Killed *C. auris*

Viable *C. auris* is not recovered in culture from bleach, ethanol and heat treatment. However, the effective environmental disinfects for *C. auris* are Clorox-based products (20). Generally recommended cleaning practices of the environmental surfaces are wiping with 10% bleach followed by 70% ethanol. Since decontamination of linen products of the healthcare facilities require high temperatures, we also included a high temperature of 90°C. No Ct values were obtained for ITS2 cDNA of bleach-treated cells while Ct values of ethanol treated cells for 1 min to 30 min were unchanged as compared to untreated cells (Fig. 2A), or when incubated for 10 min with 70% ethanol followed by washing and then incubation for prolonged period of time (30 min to 48 h) at room temperature (Fig. 2B). RNA was also stable with heat treatment as Ct values soon after heat treatment were comparable to the Ct values of untreated cells. It remained unchanged until 24 h post-incubation with significant degradation (*P* < 0.05) at 48 h post-incubation (Fig. 2B).

**Fig. 2A.**
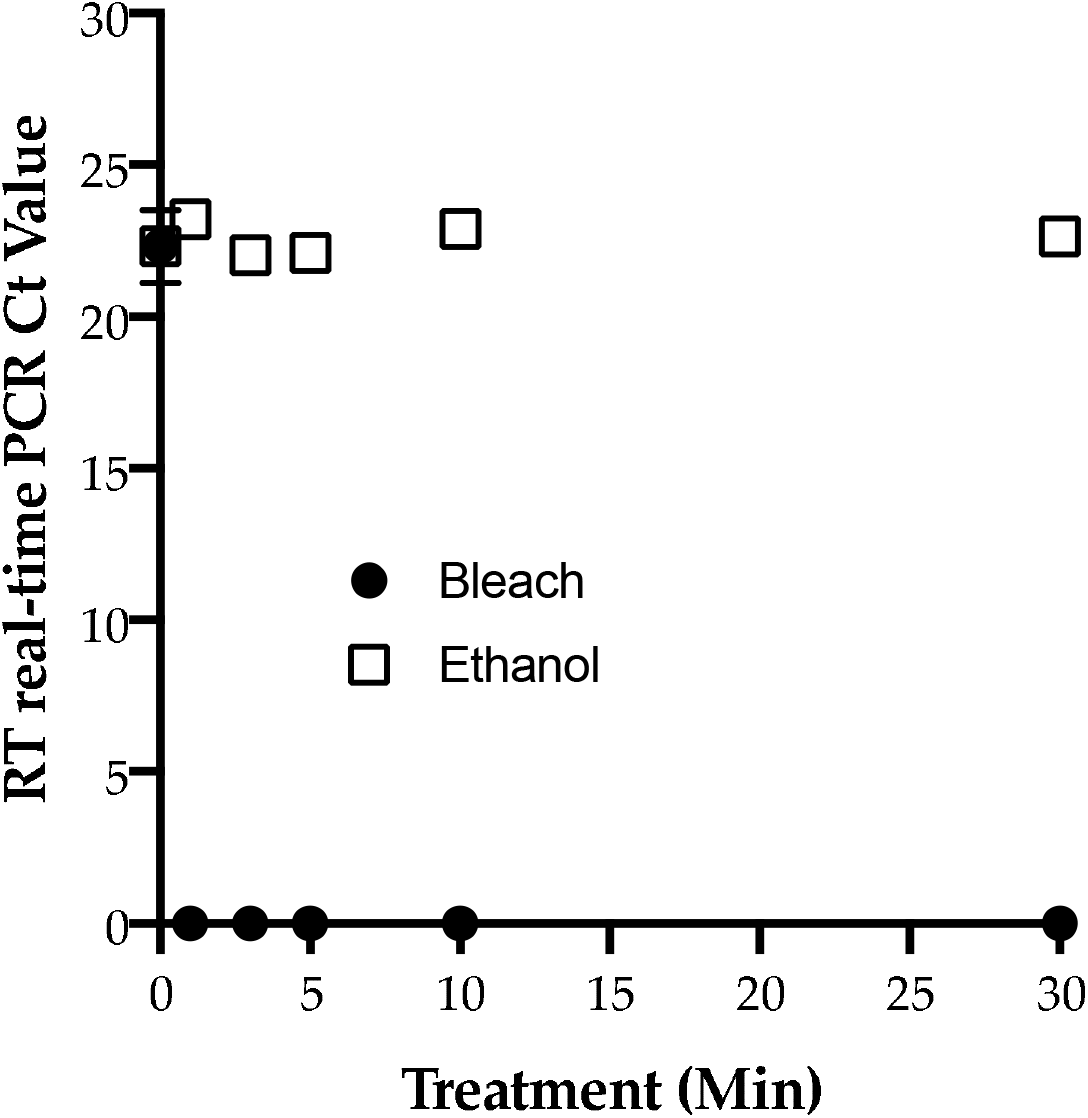
*C. auris* RNA stability following treatment with bleach and ethanol at various time points.

**Fig. 2B.**
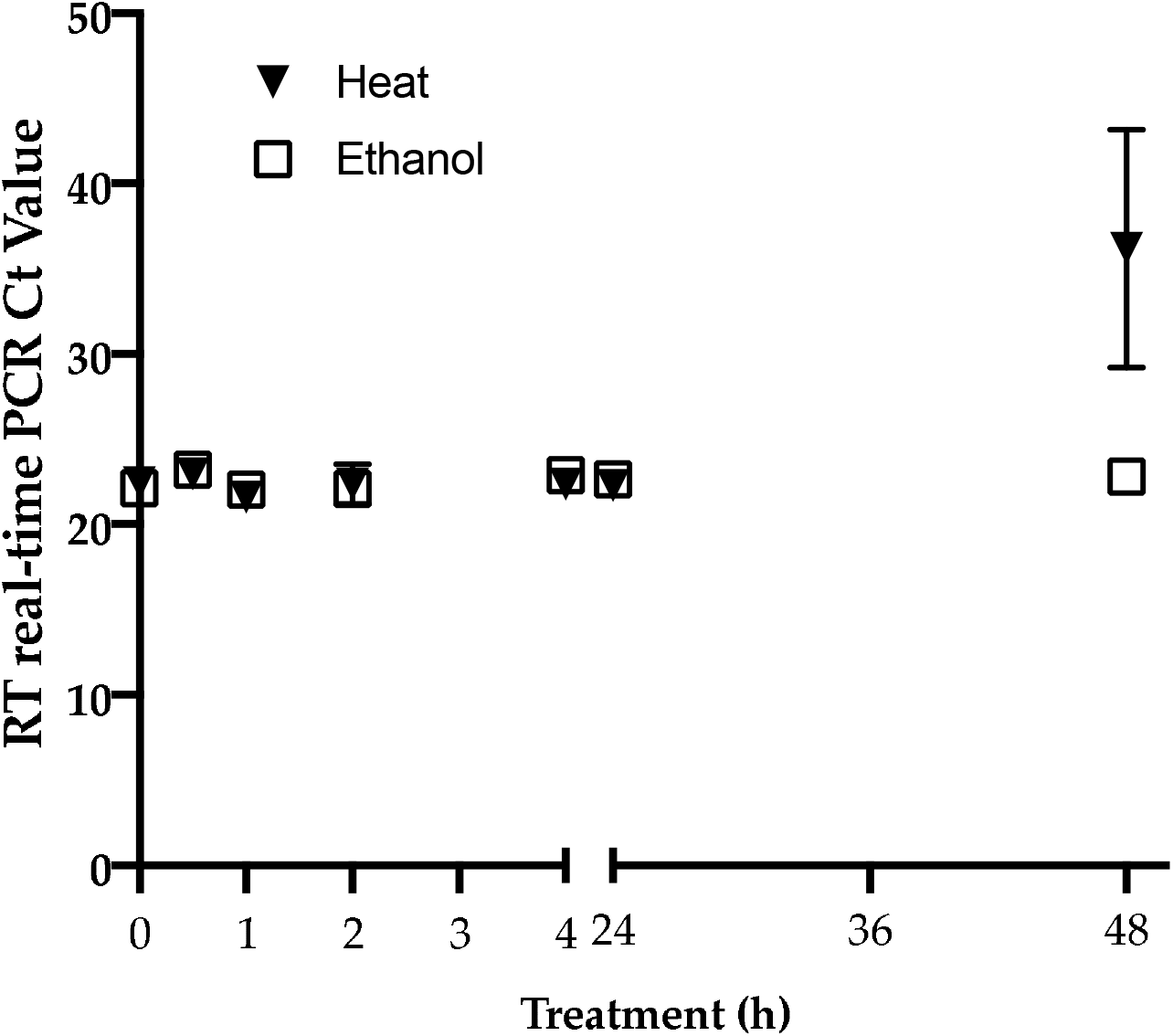
*C. auris* RNA stability following prolonged incubation at room temperature of heat and ethanol treatment.

### Retrospective Analysis of Environmental Surveillance Samples

Of 33 previously analyzed samples, 24 were positive for *C. auris* DNA by our earlier established rt-PCR assay (13) while they were all negative by culture. When these samples were subjected to RNA extraction followed by reverse transcription and real-time PCR assay, all of them were negative for ITS2 cDNA, confirming culture results (Table 2).

**Table 2.** Retrospective analysis of environmental samples from year 2020

| Sample No. | Culture | rt-PCR<br>Mean Ct<br>value | RT rt-PCR |  | Final<br>Interpretation |
| --- | --- | --- | --- | --- | --- |
|  |  |  | Ct 1 | Ct 2 |  |
| 1 | NEGATIVE | 35.23 | 0.0 | 0.0 | NEGATIVE |
| 2 | NEGATIVE | 33.71 | 0.0 | 0.0 | NEGATIVE |
| 3 | NEGATIVE | 37.26 | 0.0 | 0.0 | NEGATIVE |
| 4 | NEGATIVE | 32.17 | 0.0 | 0.0 | NEGATIVE |
| 5 | NEGATIVE | 34.38 | 0.0 | 0.0 | NEGATIVE |
| 6 | NEGATIVE | 31.62 | 0.0 | 0.0 | NEGATIVE |
| 7 | NEGATIVE | 32.65 | 0.0 | 0.0 | NEGATIVE |
| 8 | NEGATIVE | 32.14 | 0.0 | 0.0 | NEGATIVE |
| 9 | NEGATIVE | 35.58 | 0.0 | 0.0 | NEGATIVE |
| 10 | NEGATIVE | 37.08 | 0.0 | 0.0 | NEGATIVE |
| 11 | NEGATIVE | 32.62 | 0.0 | 0.0 | NEGATIVE |
| 12 | NEGATIVE | 36.32 | 0.0 | 0.0 | NEGATIVE |
| 13 | NEGATIVE | 33.54 | 0.0 | 0.0 | NEGATIVE |
| 14 | NEGATIVE | 35.51 | 0.0 | 0.0 | NEGATIVE |
| 15 | NEGATIVE | 34.33 | 0.0 | 0.0 | NEGATIVE |
| 16 | NEGATIVE | 28.43 | 0.0 | 0.0 | NEGATIVE |
| 17 | NEGATIVE | 35.73 | 0.0 | 0.0 | NEGATIVE |
| 18 | NEGATIVE | 36.26 | 0.0 | 0.0 | NEGATIVE |
| 19 | NEGATIVE | 36.38 | 0.0 | 0.0 | NEGATIVE |
| 20 | NEGATIVE | 0.0 | 0.0 | 0.0 | NEGATIVE |
| 21 | NEGATIVE | 0.0 | 0.0 | 0.0 | NEGATIVE |
| 22 | NEGATIVE | 0.0 | 0.0 | 0.0 | NEGATIVE |
| 23 | NEGATIVE | 0.0 | 0.0 | 0.0 | NEGATIVE |
| 24 | NEGATIVE | 0.0 | 0.0 | 0.0 | NEGATIVE |
| 25 | NEGATIVE | 0.0 | 0.0 | 0.0 | NEGATIVE |
| 26 | NEGATIVE | 0.0 | 0.0 | 0.0 | NEGATIVE |
| 27 | NEGATIVE | 0.0 | 0.0 | 0.0 | NEGATIVE |
| 28 | NEGATIVE | 0.0 | 0.0 | 0.0 | NEGATIVE |
| 29 | NEGATIVE | 32.20 | 0.0 | 0.0 | NEGATIVE |
| 30 | NEGATIVE | 35.4 | 0.0 | 0.0 | NEGATIVE |
| 31 | NEGATIVE | 37.07 | 0.0 | 0.0 | NEGATIVE |
| 32 | NEGATIVE | 35.21 | 0.0 | 0.0 | NEGATIVE |
| 33 | NEGATIVE | 34.37 | 0.0 | 0.0 | NEGATIVE |
| NEC | NA | 0.0 | 0.0 | 0.0 | NEGATIVE |
| XENO | NA | NA | 31.57 | 31.56 | POSITIVE |
| NTC | NA | 0.0 | 0.0 | 0.0 | NEGATIVE |
| PEC | POSITIVE | 29.46 | 29.32 | 28.46 | POSITIVE |

## DISCUSSION

We developed the new RT real-time PCR assay based on ribosomal RNA (ITS2 cDNA) as a cell viability marker, as RNA is less stable than DNA after cellular death (18). The selection of ITS RNA allowed us to recycle proven primers and probes from a previous *C. auris* real-time PCR DNA assay (13). The numerous replicates obtained for RT real-time PCR assay sensitivity and the small standard error among these replicates proved that the assay was highly robust and reproducible. The assay’s detection limit was ten colony-forming units of *C. auris* per PCR reaction, confirming the high sensitivity. The RT real-time PCR assay was highly specific as it did not cross react with closely or distantly related *Candida* spp.

*C. auris* causes prolonged contamination of the healthcare environment (12). In the absence of a known ecological niche, the presence of *C. auris* on inanimate objects in the healthcare setting could serve as source for new infection (2, 4, 5). Therefore, infection control requires extensive decontamination of floors and objects in the vicinity of the *C. auris*-positive patients (21). The preferred decontamination agents have sodium hypochlorite (bleach) as an active ingredient (https://www.cdc.gov/fungal/candida-auris/c-auris-infection-control.html). Until now, we and other have heavily relied on the rapid DNA-based real time PCR assays to assess the effectiveness of the decontamination procedures since culture results take much longer (4-14 days). However, there is a low concordance between *C. auris* DNA-positive and culture-positive results for environmental samples (5). Apparently, the results of the real-time PCR assay are skewed by the presence of *C. auris* DNA from live, dead, and growth-defective *C. auris* (5). Thus, laboratories that support ongoing decontamination efforts need a rapid test for the environmental samples that shows good concordance with the culture results.

In the present study, decontamination was simulated by treatment of *C. auris* with 10% bleach, 70% ethanol and 90°C heat. The rationale for these three conditions was: (1) the products with the active ingredient, sodium hypochlorite (bleach) (https://www.epa.gov/pesticide-registration/selected-epa-registered-disinfectants#candida-auris), have proved to be effective in decontamination of *C. auris* from health care environment, (2) the use of 70% ethanol following bleach treatment is standard practice to remove residual bleach from inanimate objects as part of safety protocols for staff and for the sustainability and durability of healthcare objects, and (3) heat-killing at high temperature is a requirement for cleaning of linens of the healthcare facilities facing *C. auris* outbreak (https://www.cdc.gov/fungal/candida-auris/c-auris-infection-control.html). Bleach treatment was effective, as no *C. auris* ITS2 cDNA was detected by RT real-time PCR from as little as 1 min post-treatment. Surprisingly, *C. auris* RNA was stable in the ethanol-killed cells, but it was significantly degraded in heat-killed cells after 48 h post cell death. Since the incubation conditions after heat or ethanol treatment were identical, the difference observed might be related to the effect of heat or ethanol treatment on RNA breakdown in dead cells. Results similar to our observations were reported for both eukaryotic (*Saccharomyces cerevisiae*) and prokaryotic (*E. coli*) organisms (17, 18). The excellent concordance observed between live- and bleach-treated *C. auris* and RT real-time PCR Ct values for spiked environmental samples affirms the current choice of sodium hypochlorite (bleach) products for the decontamination of the healthcare environment (11). Furthermore, the retrospective analyses of 33 environmental samples showed an excellent correlation between negative RT real-time PCR values to those of negative culture results.

There are some obvious limitations to our study. As RNA extraction is more labor-intensive than crude DNA extraction, the new assay will not be as high-through put as our previous assays (13, 14). Quantitative PCR systems based on reverse transcription have mostly used mRNA as the template, whereas our primers were designed to amplify rRNA (*ITS2*). Detection of mRNA might be a better technology as their shelf life is shorter than RNA. Despite their potential advantages, mRNA-based approaches have proved difficult to exploit because of the complexity of the methods, the practical problems of extracting detectable levels of intact mRNA from small numbers of cells, and a lack of basic information about the significance of detecting mRNA in the stressed cells. As ITS RNA is stable in *C. auris* cells treated with heat or ethanol, our test will only work with surveillance samples obtained after recommended decontamination with sodium hypochlorite-based agents. Lastly, our retrospective evaluation of environmental samples only showed correlation between RNA and culture for the negative samples. The RNA extraction using an Epicenter kit followed by removal of residual DNA by TURBO DNase proved to be excellent for isolation of pure RNA. DNA contamination from as high as 10^6^ *C. auris* cells was almost nil to minimal level (Ct = >40.0) as determined by real-time PCR assay.

In conclusion, the new *C. auris* RT real-time PCR is a fast, direct (without culture), sensitive, and reliable technique for the detection of live *C. auris* cells from the surveillance samples of the healthcare environment. This newly developed assay can strengthen ongoing decontamination efforts for the effective control and prevention of *C. auris* in the healthcare environment.

## ACKNOWLEDGMENTS

We thank Wadsworth Center (WC) Tissue Culture & Media Cores for providing various media for culture of *Candida* spp. We also thank Dr. Kimberlee McClive-Reed for the editing of the manuscript. This work was supported partly by the funds from the WC, New York State Department of Health (NYSDOH), and Centers for the Disease Control and Prevention (CDC) grant number NU50CK000516. In addition, Bryanna Freitas is partially supported with WC Fellowship Program. The contents of this manuscript are solely the responsibility of the authors and do not necessarily represent the official views of the NYSDOH or the CDC.

## AUTHOR CONTRIBUTIONS

SC conceived the study, supervised experiments and wrote the manuscript. BF performed majority of the experiments and prepared graphs and tables. LL was instrumental in assessing various experimental and control reagents for performing RT real-time PCR assay, supervised part of BF’s work, and tabulated previously analyzed data for the retrospective study. VC contributed to the study design and edited the draft manuscript.

## LEGENDS

**Figure 1. Sensitivity of RT real-time PCR assay**. Serial 10-fold dilution of spiked *C. auris* cells in PBS-BSA and environmental sponge samples were prepared, RNA was extracted, and RT real-time PCR was run in triplicate. The assay was linear over five orders of magnitude with assay sensitivity of 10 CFU/PCR reaction.

**Figure 2 A-B. *C. auris* RNA stability following treatment with bleach, ethanol, and heat**. *Candida auris* grown in SDA for 72 h at 30°C was scrapped, suspended in PBS-BSA, and OD_530_ = 0.1 was determined. One ml aliquot of 0.1 OD (equivalent to 10^6^ CFU/ml) was treated with 10% bleach, 70% ethanol, and heat at 90°C. **(A)** Treatment with bleach and ethanol for different time points followed by RNA extraction and RT-real-time PCR assay. **(B)** Heat treatment for 1 h followed by incubation at room temperature for 0 min to 48 h; ethanol treatment for 10 min followed by washing in PBS-BSA and incubation at room temperature for 0 min to 48 h. Both sets of samples were processed for RNA extraction followed by RT real-time PCR assay. The Ct values are the average of results from three replicates. Error bars represent the standard deviation.

**Supplementary Table 1.**
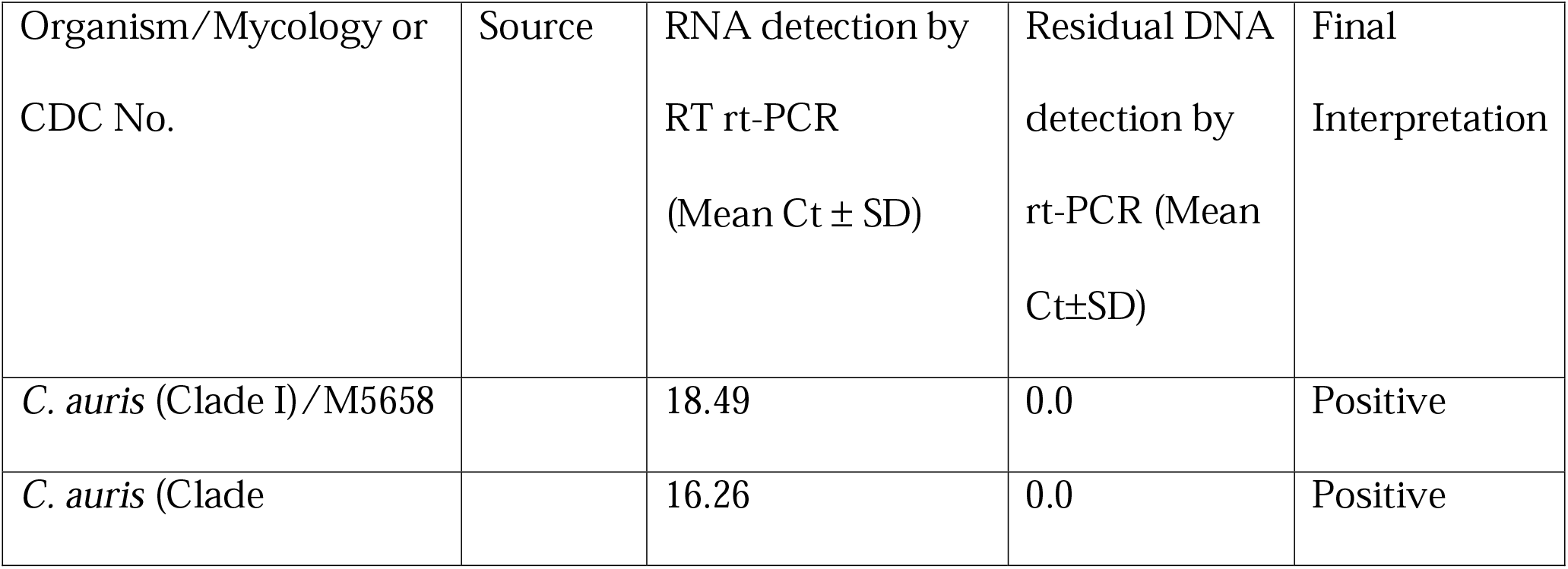
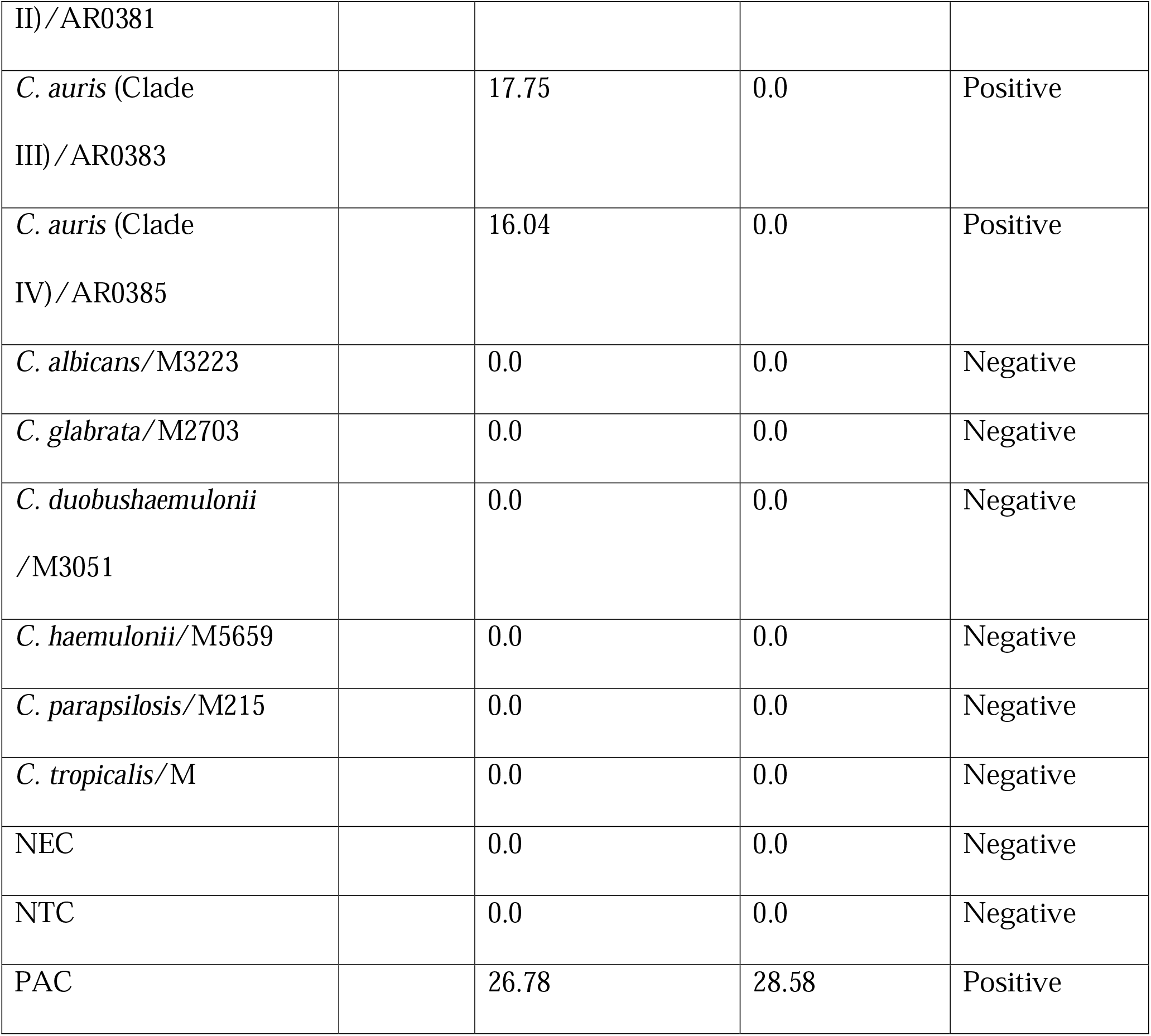
*C. auris* RT rt-PCR assay specificity

## Notes

### Competing Interest Statement

The authors have declared no competing interest.

